# Circadian multimodal physiological, ocular and behavioural data from a 40-hour forced-desynchrony protocol

**DOI:** 10.64898/2026.10.28.762537

**Authors:** Hannah S. Heinrichs, Chiara Zanoli, Salma M. Thalji, Lisa Gagelmann, Emin Kilic, Manuel Spitschan

## Abstract

We present a multimodal dataset from twelve healthy adults (aged 19–30 years, 50% female) who completed a 40-hour ultra-short sleep-wake forced desynchrony protocol which scheduled 2 h 30 min of wake-fulness in dim light and 1 h 15 min of sleep in darkness (2:1 wake:sleep ratio), with the protocol ending after the last wake period. This schedule compressed the typical 24-hour day into an ultradian 3 h 45 min cycle to decouple circadian and homeostatic influences on physiology and behaviour. During each wake interval, participants completed repeated assessments of visual psychophysics (chromatic temporal contrast sensitivity), pupillometry with silent-substitution stimuli, ocular structure (optical coherence tomography) and physiology (intraocular pressure), psychomotor vigilance, salivary hormones, and standardised questionnaires, alongside continuous actigraphy, electrocardiography, and recordings of core and peripheral temperature, blood pressure, and interstitial glucose. Two supplementary evening sessions under experimental light exposure and dim-light conditions provide comparison data on circadian phase (salivary melatonin concentration, dim-light melatonin onset). Data collected from one pilot participant and screenings of two excluded candidates are additionally included. This dataset supports reuse for investigating circadian dynamics across multiple domains beyond the retinal-function hypotheses of the original study.

## Background & Summary

### Scope of the study

Sleep and wakefulness, along with numerous downstream physiological and behavioural processes, are jointly regulated by two interacting processes^1,2^: a circadian process (“Process C”), generated by the suprachiasmatic nucleus of the hypothalamus and primarily influenced by light exposure, and a sleep homeostatic process (“Process S”), which reflects the accumulation and dissipation of sleep pressure as a function of prior time spent awake and asleep. Under habitual living conditions these two processes are tightly coupled, which makes it difficult to attribute any given physiological or behavioural outcome specifically to one process rather than the other. Forced desynchrony (FD) protocols address this by imposing a sleep-wake schedule with a period other than 24 hours; as a consequence, the circadian phase progresses independently of the sleep-wake cycle over the course of the protocol, allowing circadian and homeostatic contributions to be experimentally disentangled^3,4^.

Specifically, we used a so-called multiple-nap or “ultra-short sleep-wake” protocol^4^, a FD variant which compresses this dissociation into cycles of only a few hours, allowing more than one circadian phase to be sampled within a short overall protocol duration. Additionally, light exposure during wake periods and ambient temperature were kept constant to facilitate temporal isolation, while other daily-life aspects, such as meals or activities, were held consistent across the imposed cycle. However, this requires around-the-clock monitoring in a tightly controlled environment together with high participant compliance, making these protocols logistically demanding to run, especially if they encompass densely repeated measures. Therefore, few open datasets exist that document the full range of signals that such designs can capture.

FD protocols in humans have previously been used mainly to characterise circadian and homeostatic contributions to neurobehavioural performance^5,6^, mental health^7^ and core body temperature regulation^8^. The protocol underlying this dataset was designed primarily to investigate circadian influences on temporal contrast sensitivity^9^ and pupillary light responses^10^, examining both image-forming and non-image-forming visual functions while disentangling circadian effects from homeostatic influences.

Dissociating these influences required continuously monitoring possible confounding physiological processes throughout the 40-hour protocol, and therefore the study additionally generated dense, concurrent recordings of several environmental and physiological parameters. This Data Descriptor releases that complete underlying multimodal dataset, extending its value well beyond the retinal-function hypotheses that originally motivated data collection, and offering a resource for new hypotheses and secondary analyses within these additional physiological domains.

### Study description

The dataset presented here derives from a 40-hour ultra-short sleep-wake forced desynchrony protocol organised as an 11-cycle schedule that ended after the wake period of the final cycle. The registered study protocol^11^ specifies the full experimental setup for every recorded modality. Formal hypotheses and pre-planned confirmatory analyses were defined only for the image-forming and non-image-forming visual outcomes; the remaining modalities, including cardiovascular activity, thermoregulation, interstitial glucose, and vigilance, were documented methodologically as part of the protocol, but without pre-registered hypotheses. The confirmatory psychophysical (temporal contrast sensitivity, TCS) and pupillometric outcomes have already been analysed and reported separately by Heinrichs and Spitschan (2026)^9,10^.

This Data Descriptor accompanies those publications by releasing the complete underlying raw multimodal dataset on which they were based, spanning ocular and visual measures, cardiovascular activity, core and peripheral thermoregulation, metabolic measures, actigraphy, and self-reported data. Specifically, it reports a multimodal dataset obtained by the dense concurrent capture of these measures over three experimental sessions: an evening session with experimental light exposure (01_expsession), a dim-light evening session (02_expsession), and the 40-hour forced desynchrony protocol (03_expsession).

In addition to this main protocol, the dataset includes two supplementary recordings. The first is a pilot session in which a participant completed a dim-light evening, an adaptation night, and the full forced desynchrony protocol (approximately 53 h in total) before official data collection had formally begun, and while archiving procedures were still being finalised. As a result, pilot data are incomplete for several modalities; however, environmental control was already fully in place and the psychophysical testing was successful, hence the pilot TCS data were included in the analyses by Heinrichs and Spitschan (2026)^9,10^.

The second supplementary session is a diurnal control (DC) sub-experiment, conducted with the pilot participant 113 and the main-study participant 202, who were reinvited in July/August 2025. This session ran from 08:00 to 20:30 and consisted of 10 blocks of 1 h 15 min each; each block corresponded to one third of a full 3 h 45 min FD cycle, with alternating constrained-pupil and free-viewing conditions. No homeostatic manipulation was imposed (i.e., no scheduled sleep), and only TCS psychophysics and pupillometry (recorded with a Pupil Core eye tracker, Pupil Labs) were collected; this sub-experiment is described in the supplementary material to Heinrichs and Spitschan (2026, preprint)^9,10^.

### Reuse value

Beyond its primary relevance to retinal and visual function, this dataset holds broader reuse value, extending well beyond its original design. Each of the recorded physiological domains (cardiovascular activity, core and peripheral thermoregulation, interstitial glucose, and psychomotor vigilance) is known to be under circadian and/or homeostatic control, yet these domains are rarely co-recorded within a single ultra-short sleep-wake schedule. This concurrent, densely sampled, multimodal structure opens several concrete opportunities for secondary use, further specified in Usage Notes. For example, the dataset can support the evaluation and benchmarking of algorithms for circadian-phase estimation, methodological work on aligning and synchronising multimodal time series recorded at different sampling rates and cadences, and studying within-person coupling among temperature, interstitial glucose, vigilance, and cardiovascular measures across a circadian cycle. The intensive, repeated-measures design also provides a testing space for developing missing-data handling methods suited to dense intensive longitudinal experiments, given the device-level gaps documented in this Data Descriptor. Finally, because the protocol imposed alternating periods of scheduled wake and scheduled sleep, the dataset allows examination of the effects of scheduled sleep-wake state on these physiological measures. Users should note, however, that the sleep-wake labels reflect the imposed protocol condition rather than physiologically verified sleep staging, as no polysomnography was performed. Polysomnography was trialled during a few sleep-wake cycles of the pilot session but was ultimately deemed unfeasible given the tight experimental schedule and the burden it would have placed on participants. Salivary melatonin concentration data from this study have already been used as a validation dataset for the dlmoR dim-light melatonin onset (DLMO) hockey-stick algorithm^12^.

## Methods

### Study design

#### Study sample

The study was designed to obtain a final sample of 12 participants; recruitment was therefore conducted on a rolling basis, with candidates identified sequentially by the numbers 200–213 as they completed an online screening. Following an additional in-person screening, two candidates were excluded: participant 203, because impaired colour vision was identified using Hardy-Rand-Rittler (HRR) plates^13^, and participant 208, because they were unable to keep their eyes open during pupillometry screening, preventing reliable measurements. Recruitment continued until 12 participants had completed the full protocol. Data from the excluded participants are therefore not present in this dataset, with the exception of the screening recordings, which are included for both and retained in the screening directory. The remaining 12 participants (IDs 200–202, 204–207, 209–213) completed the full protocol with no post-enrolment dropout, constituting the final effective study sample. Each participant underwent the full within-participant protocol over a period of approximately three weeks. An additional pilot participant (ID 113) completed a reduced protocol comprising the dim-light session, the adaptation night, and the forced desynchrony session only, without the experimental light-exposure session.

#### Recruitment

Participants were recruited using flyers, posters, mailing lists, word-of-mouth, and online advertisements. Candidates completed an online screening survey, from which eligibility was determined automatically. Eligible candidates were invited to participate; all others were removed from the participant pool.

In exchange for their time, participants who also successfully completed the in-lab screening received a maximum total remuneration of EUR 600, obtained as follows: participants received EUR 30 for each evening session, EUR 20 for each adaptation night, and EUR 400 for the 40-hour forced desynchrony session; additionally, participants received EUR 20 upon returning the actigraph.

#### Inclusion and exclusion

Invitations to the in-lab screening were based on participant-reported information provided during the online screening. Consequently, these criteria were not reassessed in the laboratory, where only the remaining technical criteria were evaluated. Both sets of information are provided in screening/screening_online/ and screening/screening_lab/.

The rationale for the inclusion and exclusion criteria is described in the published protocol^11^. However, two deviations from that version should be noted: the inclusion criterion of “prioritisation of high sleep and circadian regularity” was not applied, and body mass index was calculated using weight information from the online screening. Weights measured during the in-lab screening (where participants were weighed in everyday clothes, without shoes) were instead used to calculate participants’ caloric requirements for meal preparation, not as an exclusion criterion. Consequently, in some cases there are unresolved discrepancies between the two sets of screening data, e.g., due to fluctuations in participants’ weight, but we do not believe these have any consequences for the dataset.

The list of inclusion and exclusion criteria that were ultimately applied is shown in Tables 1 and 2.

**Table 1:**
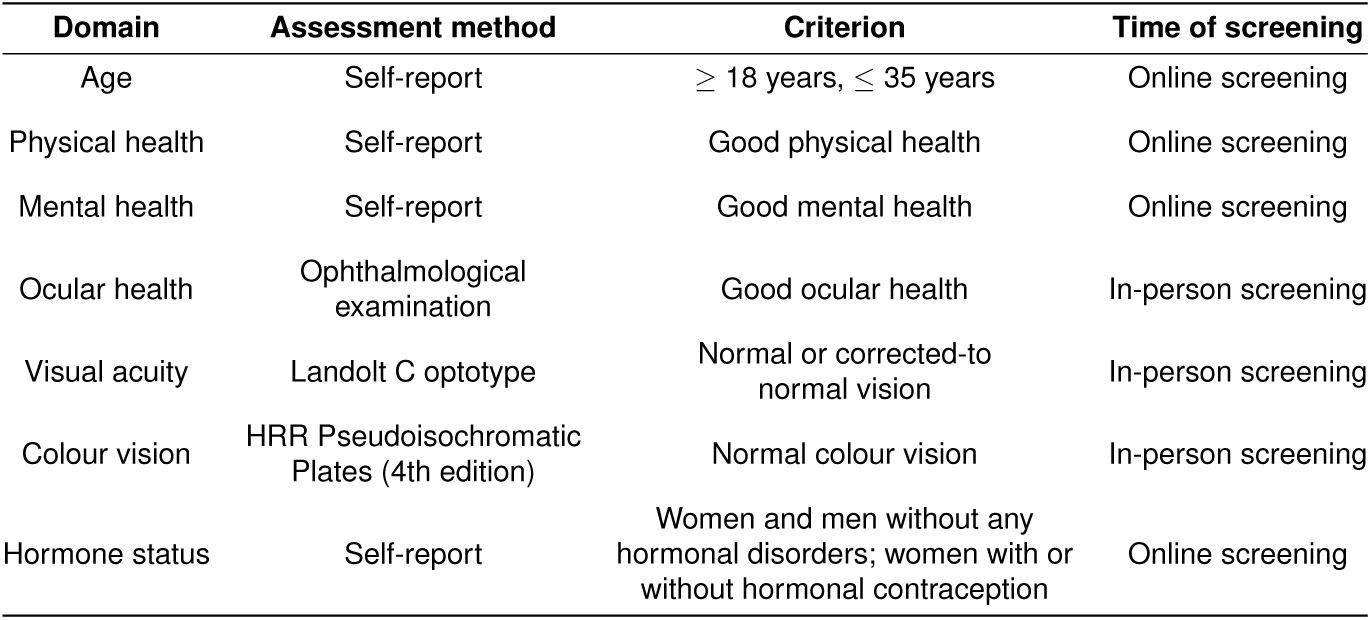
Inclusion criteria. Abbreviations: *HRR, Hardy-Rand-Rittler*^13^.

| Domain | Assessment method | Criterion | Time of screening |
|---|---|---|---|
| Age | Self-report | ≥ 18 years, ≤ 35 years | Online screening |
| Physical health | Self-report | Good physical health | Online screening |
| Mental health | Self-report | Good mental health | Online screening |
| Ocular health | Ophthalmological examination | Good ocular health | In-person screening |
| Visual acuity | Landolt C optotype | Normal or corrected-to normal vision | In-person screening |
| Colour vision | HRR Pseudoisochromatic Plates (4th edition) | Normal colour vision | In-person screening |
| Hormone status | Self-report | Women and men without any hormonal disorders; women with or without hormonal contraception | Online screening |

**Table 2:** Exclusion criteria. Abbreviations: *AUDIT, Alcohol Use Disorders Identification Test; MCTQ, Munich ChronoType Questionnaire; MSFsc, mid-sleep on free days corrected for sleep debt on weekdays; AMP, amphetamine; BZD, benzodiazepines; COC, cocaine; MOR, morphine; OPI, opiates; THC, tetrahydrocannabinol; LE, experimental light exposure; FD, forced desynchrony*.

| Domain | Assessment method | Criterion | Time of screening |
|---|---|---|---|
| Body mass index (BMI) | Calculated based on body weight and height self-measured by participants | <18.5 or >25 | Online screening |
| Substance abuse | Self-report, AUDIT | AUDIT >7 | Online screening |
| Depressive symptoms | Self-report | Reported presence | In-lab screening |
| History of anxiety disorder | Self-report | Reported presence | In-lab screening |
| Extreme chronotype | Self-report, MCTQ (MSFsc) | <2:00 or >5:30 | Online screening |
| Medication use | Self-report | Any use of medication other than hormonal contraception | Online and in-lab screening |
| Smoking | Self-report | Habitual smoking | Online screening |
| Photosensitive epilepsy | Self-report | Diagnosis of epilepsy | Online and in-lab screening |
| Shift work | Self-report | No shift work in the past 3 months | Online screening |
| Transmeridian travel | Self-report | No inter-time zone travel >2 time zones in the past 3 months | Online screening |
| Pregnancy | Self-report | Reported presence | Online screening |
| Endocrine alterations | Self-report | Reported presence | Online screening |
| Drug use | AMP, BZD, COC, MOR/OPI, THC panel | Urine test<br>positive result | LE session and FD session |
| Alcohol use | Breathalyser | >0.0 % | LE session and FD session |

#### Study protocol

The complete study protocol, including not only the experimental sessions themselves but also the screening session and stabilisation phases, spanned over a period of approximately three weeks, structured as follows:

1. An initial in-person screening;
2. A circadian stabilisation phase lasting 1 week;
3. A first experimental light-exposure (LE) session lasting 5 h in the evening (a dim-light environment supplemented only by light emitted from the visual-testing devices, not a bright-light intervention – see Rationale for the sessions), extending 1 hour past habitual bedtime and followed by an 8 h adaptation night;
4. Another week of circadian stabilisation;
5. A second dim-light (DL) session lasting 4 h in the evening, ending at habitual bedtime and followed by another 8 h adaptation night;
6. A third experimental session, starting around 1 h after the participants woke up from the second adaptation night, and lasting 40 h under an 11-cycle multiple-naps forced desynchrony schedule (3 h 45 min per cycle: 2 h 30 min awake followed by 1 h 15 min asleep), with the protocol ending after the wake period of the final cycle;
7. A final circadian stabilisation week.

During the initial in-person screening, sleep, wake and meal times, as well as the total number of meals, were established according to the habitual bedtime of each participant, and are summarised in Table 4. Participants were then required to strictly adhere to this schedule throughout all three stabilisation phases, with deviations of no more than *±* 30 minutes from the agreed times.

The first two stabilisation periods immediately preceded each corresponding session, with no break before starting the experiment. Additionally, the dim-light evening was followed directly by the forced desynchrony protocol, which started approximately 1 hour after participants were woken up by the experimenters from the preceding adaptation night; during this time, they were provided with all the necessary devices and related instructions. In contrast, the interval between the LE and the second stabilisation phase varied among participants and is documented in the 20260423_session-info.xlsx metadata file. The reasons for this variability, as well as for any postponement of the LE session, were either participant noncompliance, which required the experimenter to reschedule and restart the circadian stabilisation phase before DL and FD, or the need to align the experimental timeline with the mid-follicular phase of naturally cycling female participants^11^. The entire study timeline, including this variable inter-session interval, is shown in Figure 1.

**Figure 1:**
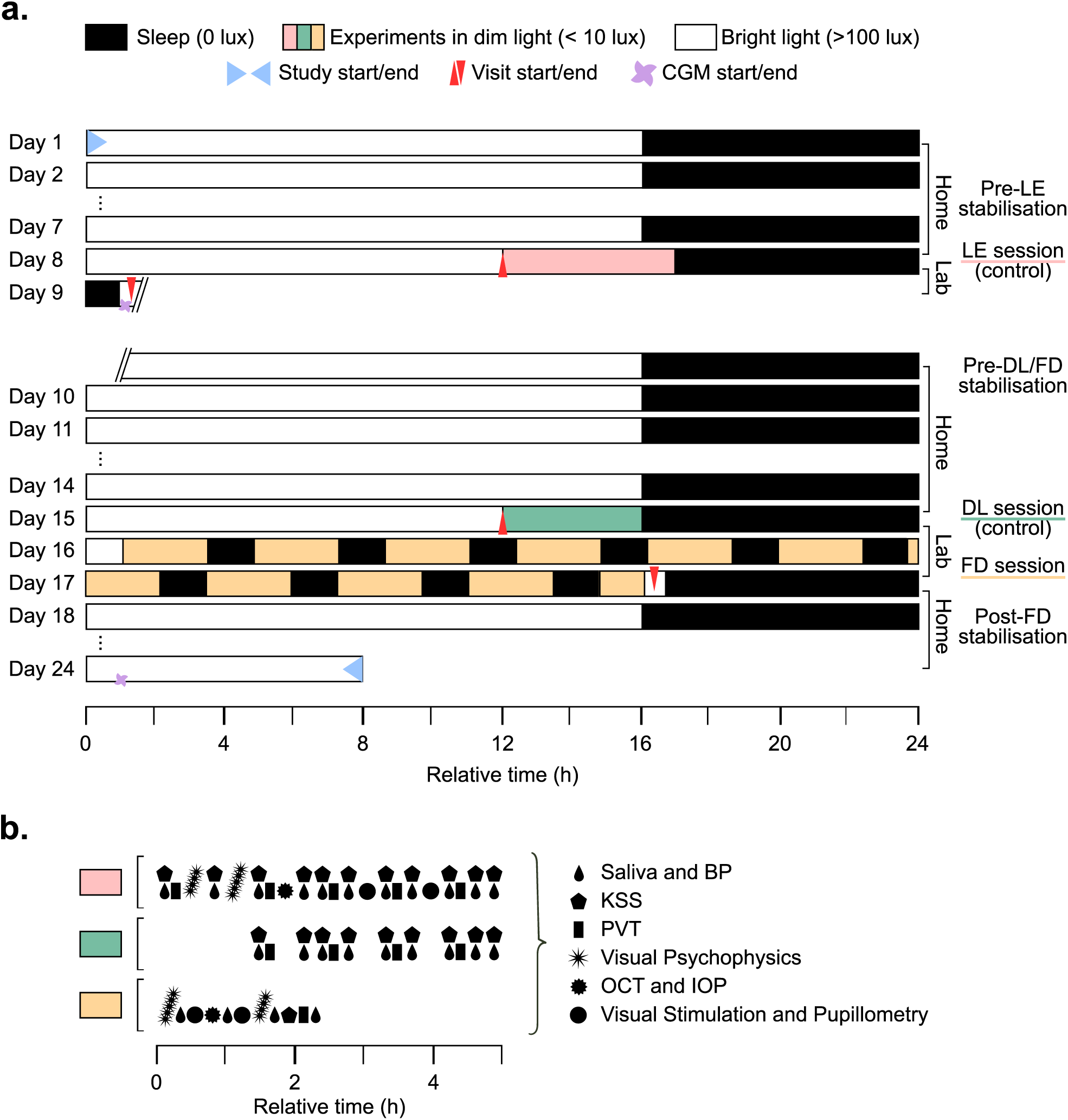
**a.** Study timeline with circadian stabilisation phases and experimental sessions; the break between Day 9 and Day 10 indicates the variable inter-session interval used to accommodate the menstrual phases of naturally cycling female participants. **b.** Details of the order and approximate timing of the different recording modalities in each experimental session. The pink and green intervals refer to the entire LE and DL sessions, respectively, whereas the yellow interval indicates one representative wake-sleep cycle of the FD session. Abbreviations: *BP, blood pressure; IOP, intraocular pressure; OCT, optical coherence tomography; PVT, psychomotor vigilance task; Visual Stim, visual stimulation*.

**Figure 2:**
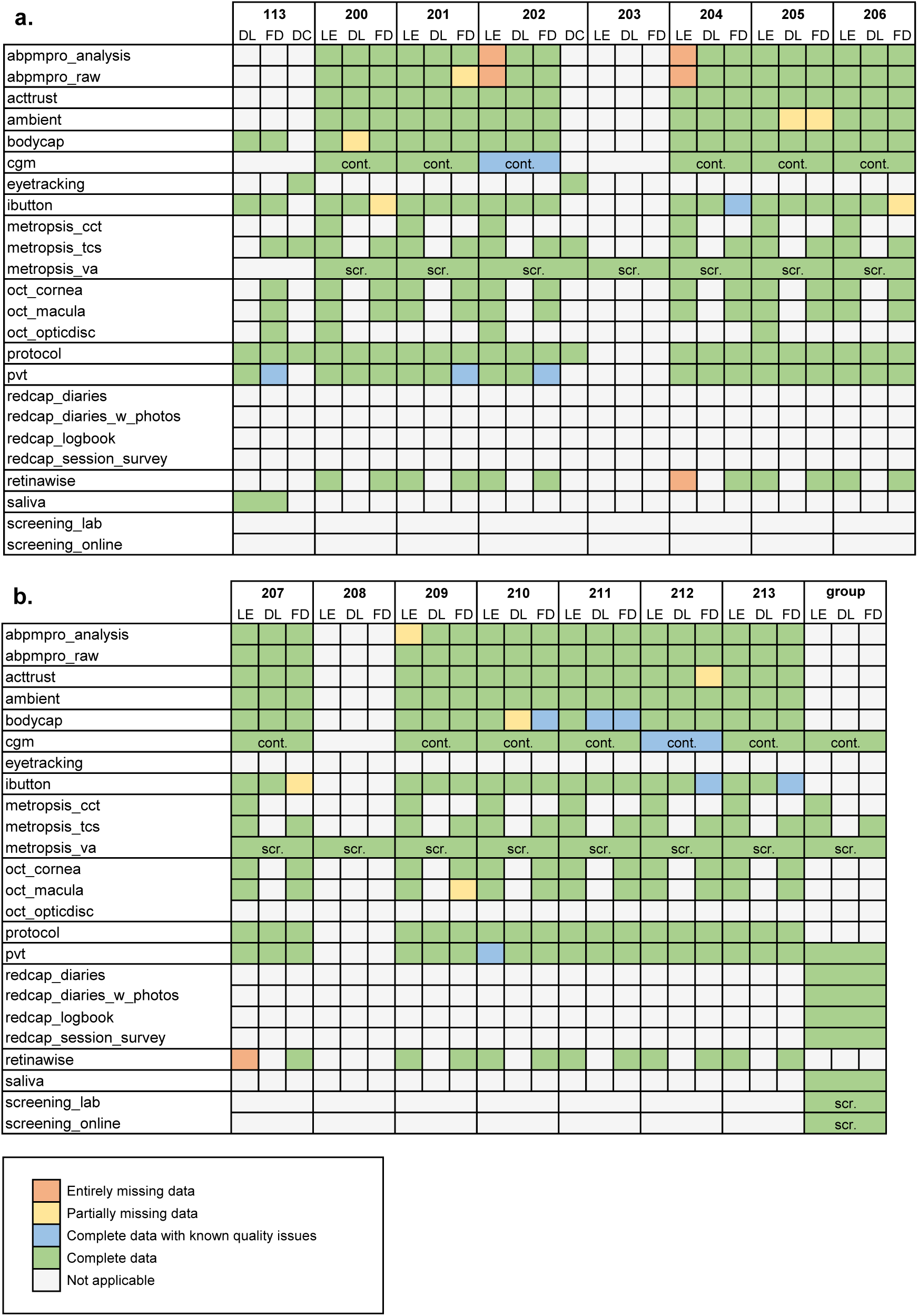
Data completeness summary across participants, modality and acquisition sessions. **a.** Pilot 113 and participants 200 to 206. **b.** Participants 207 to 213 and modalities recorded at group-level.

#### Rationale for the sessions

The LE and DL sessions were designed to assess whether light emitted by the devices used for visual experiments affected circadian and homeostatic proxies, such as vigilance, subjective sleepiness, and evening melatonin secretion. Both sessions were conducted under a *<* 10 lux light environment, with the difference that, during the LE session, the experimenters also administered the full battery of light-emitting measures: Metropsis psychophysics, retinaWISE pupillometry, and ocular imaging. By contrast, during the DL session, these measures were withheld to preserve the dim-light condition throughout the whole session, and only non-light-emitting measures were collected: salivary melatonin (for DLMO), core body temperature (CBT), distal and proximal skin temperature, and psychomotor vigilance task (PVT) performance. Comparing these two baseline sessions allowed us to exclude perturbative responses to light that could mask circadian readouts (see Technical Validation).

The 40-hour multiple-naps forced desynchrony session was instead designed to impose a highly compressed behavioural day-night cycle, thereby uncoupling the imposed sleep-wake schedule from endogenous circadian phase and allowing circadian influences to be examined separately from homeostatic influences. Furthermore, the design was intended to preserve the usual sequence of a typical waking day, alternating scheduled experimental blocks with after-work free time. FD was organised as an 11-cycle schedule, with each 3 h 45 min cycle comprising 2 h 30 min awake followed by 1 h 15 min asleep (a 2:1 wake-sleep ratio). The in-laboratory protocol ended after the wake period of the eleventh cycle; participants were then allowed to leave and complete the final sleep period at home, where it was not recorded. Isocaloric micro-meals were provided two to four times per wake interval, mirroring each participant’s habitual breakfast, lunch, dinner, and snack pattern. In the sole case of participant 207, who habitually ate five times per day (as stated in Table 4), the number of provided meals was reduced to 4 before stabilisation, due to logistical constraints. Meals were timed identically across cycles to control for metabolic and homeostatic confounds, and the caloric content of each meal was based on participants’ caloric requirements, calculated from the in-lab screening information: the participant weight measured during that screening was used to calculate the estimated caloric requirement for sedentary activity, which was then divided among the micro-meals. Within each wake interval, participants completed a fixed-order battery of assessments, including psychophysics, pupillometry, PVT, questionnaires, blood pressure measurements and saliva sampling, followed by a micro-meal. Core body temperature, skin temperature, glucose, ECG and actigraphy were recorded continuously throughout the protocol, across both wake and sleep periods. During the whole session, participants remained in a temperature-controlled room (22–23 °C, as shown in ambient/^14^ and Figure 3) under constant dim light (*<* 10 lux, outside of testing periods and except for dark sleep intervals), with no time cues available.

**Figure 3:**
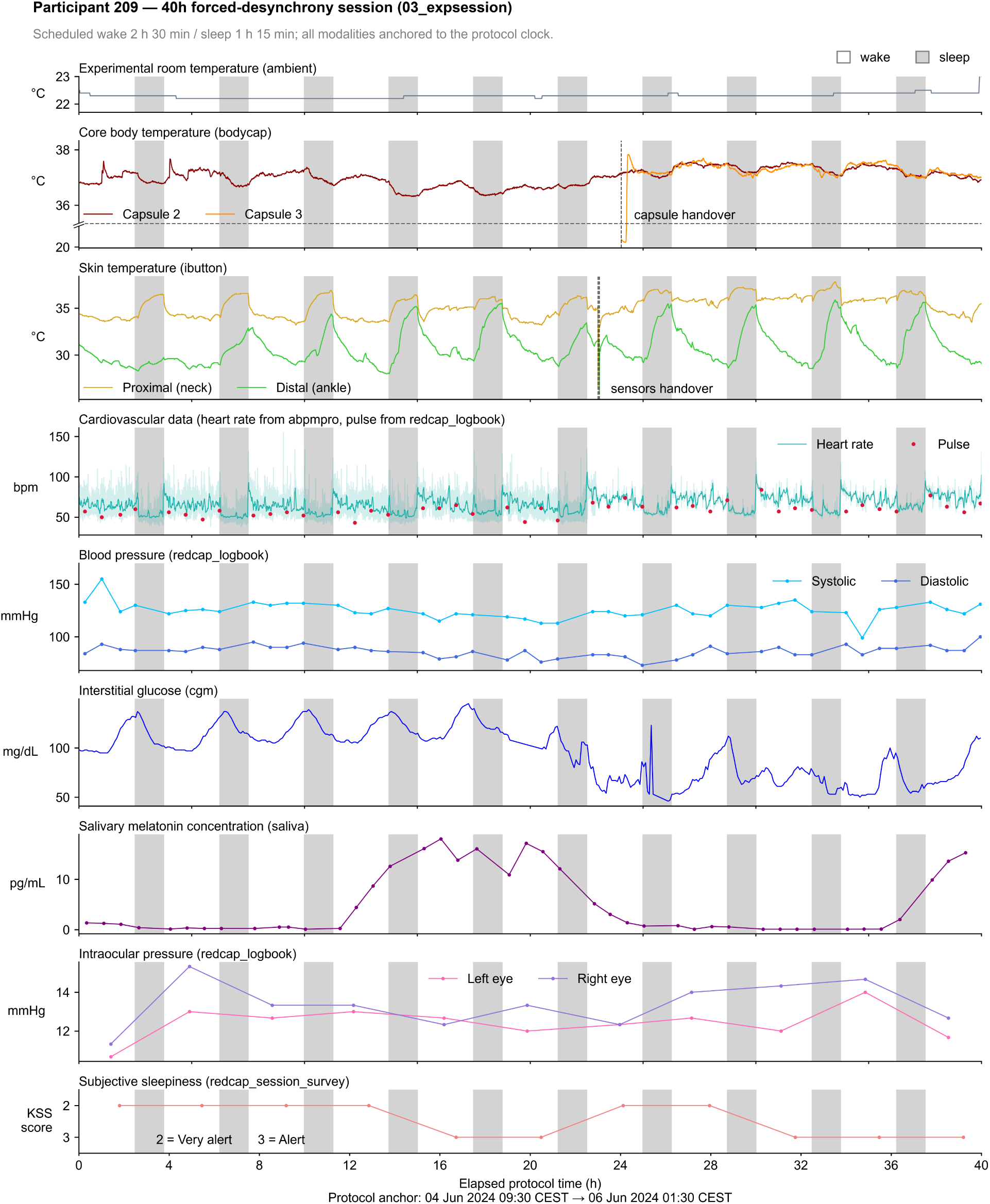
Visualisation of a selection of modalities recorded during the third experimental session (FD) of participant 209. The image depicts source data plotted over time and aligned to the FD session start. Given the high sampling density, heart rate data (measured with ABPMpro) are shown after 30-second resampling for visual clarity; the original raw data are retained in the background as a shaded trace. The background white and grey stripes indicate the imposed wake and sleep intervals, respectively. Abbreviations: *KSS, Karolinska Sleepiness Scale*.

A further diurnal control (DC) session, stored under 04_expsession, was conducted for the pilot participant and participant 202 only, in July/August 2025, after the main study period had ended. This session ran from 08:00 to 20:30 and comprised 10 75-minute blocks, with the objective of obtaining psychophysical data serving as a negative control. Unlike the FD session, it included no scheduled sleep manipulation: participants remained awake throughout the session, and a normal lunch was provided without caloric calculation, while the controlled dim-light and temperature environment was maintained as in the FD session. Blocks alternated between a free-viewing condition without any apparatus, and a constrained-pupil condition. During the latter, the dominant eye viewed through a pupil-relay system restricting pupil diameter to 1.5 mm, while the non-dominant eye was patched. Each block included temporal contrast sensitivity (TCS) testing and pupillometry, where pupil diameter was recorded via a wearable Pupil Core eye tracker, independently of the psychophysics measures.

### Data acquisition

#### Devices

Instruments used to collect each modality are summarised in Table 3, which lists the device name and manufacturer, the outcome variable, the placement and the sampling rate for each measure; full technical specifications, serial numbers, certification details, and manufacturer links are instead provided in the devices.yaml metadata file on the published repository.

**Table 3:** Summary of principal characteristics of measurement devices, matched by modality.

| **Folder name** | **Device Name** | **Software<br>(if applicable)** | **Manufacturer** | **Primary<br>Outcome/s** | **Placement** | **Sampling<br>Frequency** |
|---|---|---|---|---|---|---|
| **abpmpro** | ABPMpro | — | SOMNOmedics,<br>Randersacker, DE | Blood pressure,<br>pulse, ECG | Upper arm,<br>chest and ribs | 256 Hz, 30 s |
| **acttrust** | ActTrust2 | ActStudio 2.1.2 | Condor Instruments,<br>São Paulo, BR | Activity,<br>light exposure | Non-dominant<br>wrist | 25 Hz,<br>aggregated into<br>1-min epochs |
| **ambient** | SwitchBot<br>Thermometer and<br>Hygrometer Plus | — | Woan Technology<br>(Shenzhen) Co., Ltd.,<br>Shenzhen, CHN | Environmental<br>temperature<br>and humidity | Experimental<br>rooms | 60 s |
| **bodycap** | eCelsius<br>(+ eViewer) | eCelsius Manager | BodyCAP, Hérouville-Saint-Clair, FR | Core Body<br>Temperature | Stomach<br>(ingestible<br>capsule) | 30 s |
| **cgm** | FreeStyle Libre 3 | — | Abbott Diabetes Care<br>Ltd., Witney, UK | Interstitial<br>glucose | Back of the<br>upper left arm | 5 min |
| **eyetracking** | Pupil Core | — | Pupil Labs GmbH,<br>Berlin, DE | Pupil<br>diameter | Face | Trial-based |
| **ibutton** | iButton DS1922L | OneWireViewer | Maxim/Dallas<br>Semiconductor Corp.,<br>Dallas, Texas (USA) | Distal and proximal<br>skin temperature | Neck,<br>opposite ankle | 30 s |
| **metropsis** | Metropsis | — | Cambridge Research<br>Systems, Rochester, UK | Visual function (colour<br>vision, temporal contrast<br>sensitivity, visual acuity) | Presented<br>on screen | Trial-based |
| **oct** | Topcon 3D<br>OCT-2000 | OCT Viewer | TOPCON Corporation,<br>Tokyo, JP | Ocular structure (Optical<br>Coherence Tomography) | Both eyes | Trial-based |
| **pvt** | AMI PVT-192 | PVTCOMM 2.61 | Ambulatory Monitoring<br>Inc., Ardsley, NY (USA) | Psychomotor<br>vigilance | Dominant hand | Event-based |
| **redcap_logbook** | Auto Non-Contact<br>Tonometer<br>NT-2000 | Manually<br>recorded<br>on REDCap | NIDEK Co., Ltd.,<br>Gamagōri, JP | Intra-ocular<br>pressure | Both eyes | Trial-based |
| **retinawise** | retinaWISE | — | Oculox Technologies<br>SA, Muzzano, CH | Pupil diameter | Dominant eye | ≥ 30 Hz |

For the protocol and saliva modalities, as well as for some types of self-reported data or data logging in redcap, no devices were used, and therefore they are not included in the device list. Nevertheless, to ensure reproducibility, we report the relevant collection details here. The surveys included in the redcap folder were collected via participants’ own electronic devices (smartphones, PCs or tablets), using the web-based REDCap platform - Research Electronic Data Capture (REDCap Consortium, Vanderbilt University, Nashville, Tennessee). Saliva samples were collected using Salivette (Sarstedt AG & Co. KG, Nümbrecht, Germany) at 30-minute intervals during the LE and DL sessions and at 45-minute intervals during wake periods of the FD session. Additionally, two samples were collected in the preparation hour between DL and FD, and logged as the first two FD samples. This resulted in a total of 11 samples per participant during the LE session, nine samples during the DL session, and 46 (2 + 44) samples during the FD session. Salivary melatonin and cortisol concentrations were subsequently determined by radioimmunoassay (RK-DSM2; limit of quantification 0.5 –50 pg/mL, detection limit 0.2 pg/mL, mean intra-assay precision: 7.9%, mean inter-assay precision 9.8%) and ELISA (CRTN-96-U; detection limit 0.17 ng/mL, limits of full quantification 0.35 – >30 ng/mL, intra-assay precision 2.9-7.4% CV, inter-assay precision 5.4-11.9% CV), respectively. Neither assay was conducted in-house; both were performed by NovoLytiX GmbH, Witterswil, Switzerland.

#### Pupillometry and psychophysics specifications

Metropsis (Cambridge Research Systems, Ltd., Rochester, UK) and retinaWISE (Oculox Technologies SA, Muzzano, Switzerland) required two custom viewing apparatuses that were built as part of the experimental setup to ensure measurement stability. Psychophysical testing used a custom chin rest to maintain stable alignment, with build details described by Heinrichs and Spitschan (2026)^9^. Pupillometry was conducted inside a purpose-built blackout tent with a chin rest, designed to exclude ambient light and stabilise head and eye position; construction details are provided by Heinrichs and Spitschan (preprint)^10^. These two instruments also required participant-specific calibration. In Metropsis, stimuli were recalibrated to each participant’s refractive error at the experimental light exposure (LE) session and once at the beginning of the forced desynchrony (FD) protocol. In retinaWISE, refractive error correction was performed before each test.

For both the pupillometry and psychophysics modalities, the dominant eye was determined via the Miles test^17^ and chosen by default as the one to be stimulated, in order to minimise participants’ fatigue. Both eyes were tracked throughout pupillometry. Data from the stimulated eye were treated as the primary outcome; any deviations from this convention are documented in the Technical Validation section.

Lastly, retinaWISE is a device that has recently been renamed by the manufacturing company after its prototype was finalised as a commercial product. It can therefore be found in the earlier literature under the name “NVBL”, including in the article describing the experimental protocol associated with this data descriptor^11^.

### Data processing

The dataset consists of raw or minimally processed exports generated directly by each instrument’s proprietary software (see Table 3) without the application of any custom processing code. Where processing did occur, it was limited to instrument-internal algorithms embedded in the manufacturer’s software:

- **acttrust** Activity counts (PIM, Proportional Integral Mode) and light exposure from the ActTrust2 were preprocessed by ActStudio, which filters the tri-axial accelerometer signal, computes the vector norm across axes, and integrates it over the 60-second epoch to produce the PIM value.
- **cgm** Glucose values from the Libre 3 were preprocessed by the onboard, factory-calibrated firmware; however, Abbott does not publicly disclose the internal details of this proprietary conversion algorithm.
- **metropsis** Visual function thresholds from Metropsis were derived using the software’s built-in adaptive staircase procedures, which estimate psychophysical thresholds directly from the sequence of stimulus-response trials, making this a form of instrument-internal processing rather than a raw measurement.
- **retinawise** Pupil traces from retinaWISE reflect the device’s internal smoothing, tracking and image-processing algorithms rather than raw frame-level data. According to the the device developers^18^, this includes OpenCV-based adaptive thresholding, blob and edge detection, and ellipse fitting to determine pupil boundaries.

For the iButton temperature sensors, an additional filtered copy of the data was created for participant 204, FD session, due to three sensors identified as malfunctioning (see Technical Validation for details). These copies retain only physiologically plausible values and are stored on GitHub^15^, together with the Python script used for filtering. On the Edmond repository^14^, the corresponding raw data were preserved for transparency.

Any other corrections to the file content (e.g. re-calculations of melatonin concentrations, duplicate recordings resulting from suspected device failure, etc.) or naming (e.g. corrections to participant and session identifiers) are documented in a specific metadata file accompanying the raw data on the published repository, and annotated directly within the affected files.

File organisation in the published repository, file renaming and completeness checks were performed manually throughout. These procedures, along with any resulting gaps in the dataset, are documented in the missing_data.yaml metadata file (see Code Availability).

### Ethical approval

The study was approved by the Medical Ethics Committee of the Technical University of Munich (approval ID 2023-369-S-CB) and conducted in accordance with the Declaration of Helsinki. All participants gave written informed consent.

## Data Records

### Dataset

#### Hosting repository

The dataset (v1.0.0) is hosted at Edmond, the Open Research Data Repository of the Max Planck Society^14^. It is distributed as one compressed archive (.zip) per modality or submodality rather than as a single monolithic file, so that individual modalities can be downloaded independently of one another; each archive preserves the internal folder layout described below when extracted. A top-level DATA_TREE.txt file and the hash file SHA256SUMS_v1.0.0.txt allow users to verify the completeness and integrity of the repository structure and of individual downloads.

#### Repository structure

Within each modality folder, data are organised by participant, in folders 200 to 202, 204 to 207 and 209 to 213. Within each participant folder, data are further organised by experimental session: 01_expsession folder for the experimental light exposure (LE) session, 02_expsession for the dim-light (DL) session, 03_expsession for the forced desynchrony (FD) session, and 04_expsession for the diurnal control (DC) session, the latter being available only for participant 202 and for the pilot participant 113. A separate screening folder, placed at the same level of the modality folders, holds data that were collected solely for eligibility assessment. This additionally includes participants 203 and 208, who were excluded after screening; their data appear only in the metropsis_va folders and in the pooled screening_lab and screening_online files, with no corresponding entries in any experimental modality.

Two further structural elements sit alongside the per-participant folders. A 113_pilot folder contains data from the pilot participant, comprising only the DL and FD sessions (no LE session was run) together with a DC session collected in parallel with participant 202. Furthermore, a group folder holds data that are not organised per participant, such as the cumulative REDCap export and the pooled continuous glucose monitoring (CGM) records. Finally, a top-level METADATA folder provides project-, dataset-, device-, participant-, and missing-data-level documentation in .yaml format, accompanied by Markdown descriptors for each modality.

#### Exceptions

A number of modalities deviate from this general folder scheme, and these exceptions are documented in dataset.yaml. In particular, acttrust contains three subfolders that do not follow the standard session-naming scheme, but are instead named pre-le, pre-fd and post-fd to better reflect their content, which mainly consists of recordings obtained during the three stabilisation phases; cgm has no session subfolders at all, reflecting its continuous 14-day wear protocol; saliva exists only within the 113_pilot and group folders, as a single cumulative file; finally, the light-emitting modalities metropsis, retinawise, and oct have no 02_expsession (DL) folder, since no light exposure took place during the dim light session.

Moreover, some folders are split into submodality folders. That is the case of: abpmpro_analysis and abpmpro_raw; metropsis_cct, metropsis_tcs and metropsis_va; oct_cornea, oct_macula and oct_opticdisc; redcap_diaries, redcap_diaries_w_photos, redcap_logbook and redcap_session_survey; screening_lab and screening_online. Within these, the data are in turn organised by participant as in the other main modality folders.

### Modalities

Device references for each modality can be found in Table 3. Here, the biological meaning and repository organisation of each modality are described:

- **abpmpro** comprises brachial blood pressure measurements acquired at regular intervals, together with heart rate and three-channel ECG recordings, stored as tabular .txt files. It is further divided into two submodalities: abpmpro_raw, which contains the placement coordinates of the device components, and abpmpro_analysis, which stores the sampled data.
- **acttrust** provides continuous wrist activity and light exposure recordings, also stored as .txt files, with accompanying .pdf reports. Most of the data collected with this modality pertain to the stabilisation phases of the experimental pipeline, and are therefore stored in three folders named pre-le, pre-fd and post-fd instead of 01_expsession, 02_expsession and 03_expsession, respectively.
- **ambient** records the environmental conditions of the experimental rooms, temperature and humidity, as .csv files.
- **bodycap** uses a temperature sensor in the form of an ingestible capsule to record core body temperature, storing it in .csv files with accompanying .pdf visualisation plots.
- **ibutton** instead provides proximal and distal skin temperature recordings, saved in .xlsx format. Proximal temperature was captured with a temperature logger attached to the participant’s neck, while the sensor for distal temperature was placed on the ankle, contralateral to the neck sensor.
- **cgm** contains 14 days of continuous interstitial glucose recordings, also stored in .xlsx files. For this modality, only two readers were used across all participants; hence two group files are available. Some participants’ data are split between the two readers.
- **eyetracking** records pupil diameter during free viewing, collected during the Metropsis temporal contrast sensitivity (TCS) task, only for the DC session and across 10 alternating blocks; data are stored as .csv files, together with nested folders of raw recordings.
- **metropsis** comprises three Metropsis-based modalities capturing different aspects of visual function, all in the form of .xls files with accompanying .pdf reports: metropsis_cct records the results of the Cambridge Colour Test, collected during the LE session only; metropsis_tcs records temporal contrast sensitivity, collected during the LE and FD sessions and, for the pilot participant and participant 202, also during the DC session; metropsis_va is an assessment of visual acuity, collected only during screening, and therefore also containing data from excluded participants 203 and 208.
- **oct** also comprises three submodalities, all obtained through optical coherence tomography (OCT) scans and stored as .pdf reports; each submodality name identifies the object of the OCT scan: cornea, macula and optic disc. oct_macula additionally includes retinal thickness measurements as .xlsx and .csv files.
- **protocol** is one of the few modalities not associated with a device, since it does not sample any participant measures, but instead provides a minute-by-minute .csv log of experimental events, used as in-session audio instructions for participants and experimenters.
- **pvt** contains psychomotor vigilance task reaction-time trials, stored in the native .RTF format.
- **redcap** refers to modalities for which REDCap questionnaires were used, either for data collection through surveys administered directly to participants, or for logging experimental measures. The former is the case for redcap_diaries_w_photos and redcap_diaries, which contain surveys administered during the stabilisation phases, respectively with and without pictures of participants’ meals, and for redcap_session_survey, which stores participants’ responses to surveys collected during the experimental sessions, including for example the subjective sleepiness measured with the Karolinska Sleepiness Scale and plotted in Figure 3 and Figure 4. redcap_logbook instead contains data logged in by experimenters, including intraocular pressure captured by the auto non-contact tonometer, blood pressure and pulse, also shown in Figure 3. Each submodality includes both the survey data and the corresponding data dictionaries, stored as .csv files.
- **retinawise** records pupil diameter, pupillary light reflex, and silent substitution responses, collected during the FD session only, as .csv files.
- The **saliva** modality provides salivary melatonin and cortisol concentrations measured by radioimmunoassay and stored in a single group .xlsx file (with the exception of pilot 113, which has its own export). This modality is likewise not linked to a device; saliva samples were collected using Salivette (Sarstedt AG & Co. KG, Germany) and subsequently analysed by NovoLytiX (NovoLytiX GmbH, Switzerland).
- Finally, **screening** contains group data from two screening sessions: the preliminary recruitment of potential participants, assessed through an online questionnaire, screening_online, and the final inperson screening, which confirmed the actual participant sample of this study, screening_lab. This modality includes responses from the two excluded participants 203 and 208, with all data stored as .csv files.

**Figure 4:**
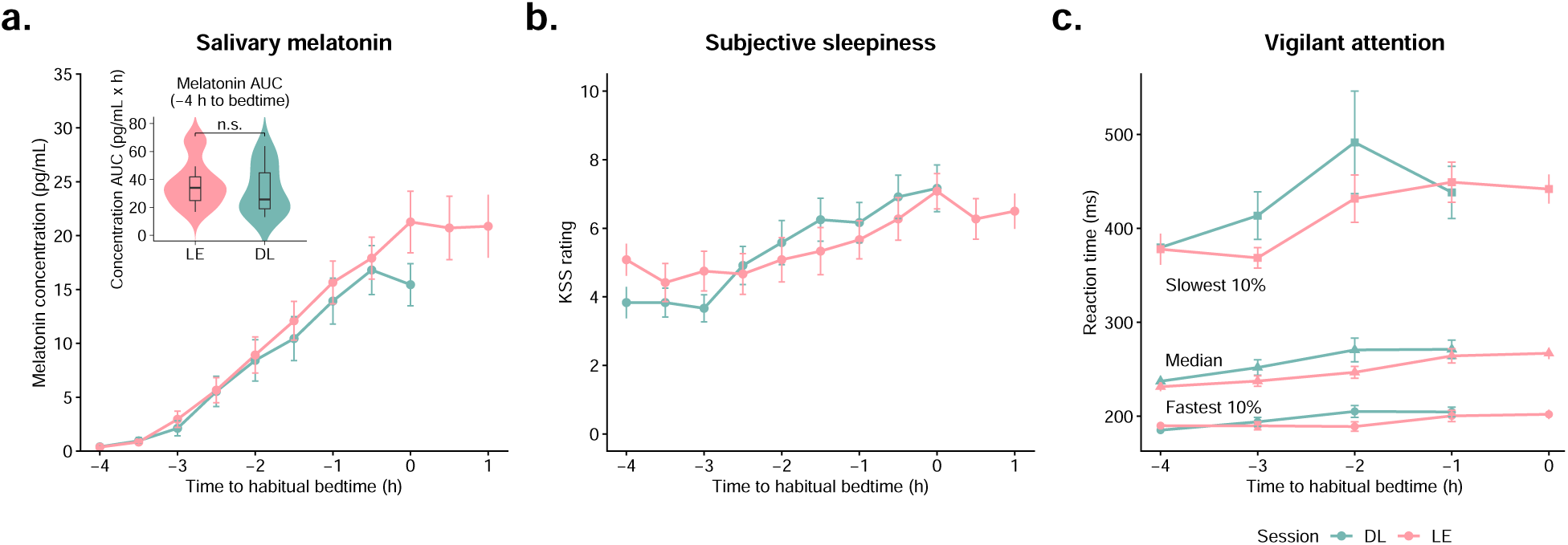
LE (pink) and DL (green) comparison over time of: **a.** salivary melatonin concentration, **b.** self-reported subjective sleepiness measured with the Karolinska Sleepiness Scale, and **c.** vigilant attention, measured by the psychomotor vigilance task (PVT). Visualisation relative to hours before habitual bedtime.

File naming conventions, header presence, and column definitions for every modality are specified in data-set.yaml and in the corresponding Markdown descriptors under METADATA/descriptors/.

### Participants

A total of 14 candidates were recruited and screened for the study, following a rolling recruitment scheme designed to yield a final sample of 12 (see Study design for recruitment details). Participants 203 and 208 were subsequently excluded because they did not meet the normal colour-vision criterion and could not complete the pupillometry assessment, respectively. The remaining 12 participants completed the full experimental protocol with no dropout over the course of the study. However, due to defective devices, export issues or protocol deviations, not all modalities were successfully collected in all sessions for each participant, and therefore completeness of the data can be assessed through Figure 2, as well as through the participants.yaml metadata file reporting full participant-level detail, including demographics, inclusion status and session history.

In addition to this study sample, data from pilot participant 113 are included in the repository. This participant took part in a separate pilot run conducted prior to the formal start of data collection, comprising only a dim-light phase, an adaptation night, and the full forced desynchrony protocol (approximately 53 h in total). As the archiving pipeline had not yet been finalised at that stage, some data modalities from this session remain incomplete, and are also listed in Figure 2.

Finally, two participants, the pilot and 202, were recalled for a diurnal control sub-experiment, which only included assessment of temporal contrast sensitivity and pupillometry (see Study description for details).

All participants appearing in this dataset, including the pilot and the subsequently excluded 203 and 208, were, at the time of enrolment, aged between 19 and 30 years (mean *±* SD: 24.4 *±* 2.9 years) and had a BMI ranging from 18.9 to 24.6 kg/m^2^ (mean *±* SD: 21.1 *±* 1.6 kg/m^2^). Half of the participants were female, either using or not using hormonal contraception; naturally cycling female participants were tested in the mid-follicular phase of their menstrual cycle, determined from the timing of menses, which is recorded in the 20260423_session-info.xlsx metadata file.

More detailed demographic characteristics of all participants are summarised in Table 4, including any health status specifications, reasons for exclusion, and the number of habitual meals selected for the experiment.

**Table 4:** Participant demographic data. Meal counts reflect participants’ habitual number of meals, except for participant 207, whose count was reduced to 4 for logistical reasons.

| Participant ID | Sex (M/F) | Age (years) | Height (cm) | Weight (kg) | BMI (kg/m<sup>2</sup>) | Number of meals | Sleep time | Wake time | Medical notes | Participant status | Reason for exclusion |
|---|---|---|---|---|---|---|---|---|---|---|---|
| **113** | M | 25 | 160 | 50 | 19.5 | n/a | 00:00 | 08:00 | — | pilot | — |
| **200** | M | 30 | 172 | 71 | 24.0 | 3 | 00:00 | 08:00 | eye surgery (LASIK) | complete | — |
| **201** | F | 22 | 170 | 60 | 20.8 | 4 | 22:00 | 06:00 | hormonal contracep-tion | complete | — |
| **202** | M | 24 | 180 | 67 | 20.7 | 2 | 22:30 | 06:30 | — | complete | — |
| **203** | F | 27 | 161 | 49 | 18.9 | 3 | 23:00 | 07:00 | naturally cycling | excluded | impaired colour vision |
| **204** | F | 22 | 162 | 55 | 21.0 | 3 | 22:00 | 05:30 | naturally cycling | complete | — |
| **205** | M | 21 | 202 | 80 | 19.6 | 3 | 23:30 | 07:15 | — | complete | — |
| **206** | M | 23 | 190 | 84 | 23.3 | 4 | 23:30 | 07:30 | — | complete | — |
| **207** | F | 19 | 165 | 55 | 20.2 | 5 | 22:30 | 07:30 | hormonal contracep-tion | complete | — |
| **208** | M | 24 | 178 | 78 | 24.6 | 4 | 01:00 | 09:00 | — | excluded | inability to complete pupillometry |
| **209** | M | 28 | 167 | 63 | 22.6 | 2 | 00:30 | 08:30 | — | complete | — |
| **210** | F | 23 | 170 | 58 | 20.1 | 4 | 23:30 | 07:00 | naturally cycling | complete | — |
| **211** | F | 25 | 157 | 52 | 21.1 | 4 | 23:45 | 08:00 | naturally cycling | complete | — |
| **212** | F | 24 | 166 | 55 | 20.0 | 4 | 23:00 | 07:00 | naturally cycling | complete | — |
| **213** | M | 29 | 175 | 60 | 19.6 | 4 | 00:00 | 08:00 | — | complete | — |
| **mean ± SD** | — | 24.4 ± 2.9 | 171.7 ± 11.7 | 62.5 ± 10.8 | 21.1 ± 1.6 | — | 23:23 ± 00:51 | 07:23 ± 00:54 | — | — | — |

### Data completeness

To support reuse of the dataset, session-level data completeness for each participant is reported in Figure 2, covering the continuous, LE, DL, FD, screening and supplementary DC sessions. However, DC-session data are limited to the pilot participant and participant 202, with no contributions from other participants, while pilot 113 completed only a subset of the modalities collected from the other participants. Therefore, for each participant and session, completeness is categorised as “Complete,” “Complete with known quality issues,” “Partially missing,” “Entirely missing,” or “Not applicable (N/A),” the latter indicating that the data were not intended to be collected according to the protocol and are therefore appropriately absent. This classification was derived by cross-referencing participants.yaml and missing_data.yaml, two metadata files that contain full specifications regarding partial rather than complete modalities, as well as reasons for the missing data.

## Technical Validation

### Visual overview

Figure 3 illustrates, for a single representative participant (209), how the released streams behave when viewed together across the full 40-hour FD protocol. Together, these traces serve as a qualitative check that the raw exports behave as expected given the study design.

Each modality shown is plotted on a common protocol-time axis anchored to the FD session start, demonstrating that timestamps across independent files can be aligned to a shared timeline despite having different sampling rates and export formats. The continuous modalities (ambient, bodycap, ibutton, abpmpro and cgm) and saliva show technically plausible traces with the expected physiological structure (e.g. visible rhythmical fluctuations). Additionally, the figure also makes visible the device handovers (the mid-protocol pill swap in the core-temperature trace and the sequential iButton sensor switch in the skin-temperature trace) and, especially in the case of non-continuously recorded data (redcap_logbook, redcap_session_survey and saliva), the sampling structure of each modality.

Raw data for pulse, blood pressure, intraocular pressure, and subjective sleepiness were extracted from the corresponding redcap export files and related dictionaries before being plotted. Interstitial glucose samples specific to participant 209 were also first derived from group-level export files.

### Instrument calibration and certification

All instruments underwent calibration or certification appropriate to their measurement type prior to data collection. The iButton temperature loggers were factory-calibrated in a NIST-traceable chamber. The Metropsis Maxwellian-view perimeter was recalibrated for refraction error both at the beginning of the LE and the FD sessions. The retinaWISE device was recalibrated once every session according to the manufacturer’s instructions. The Topcon OCT-2000 and NIDEK tonometer both carry CE/ISO certification. All calibration details and the list of certificates for each instrument are provided in devices.yaml.

### Effect of light-emitting instrumentation

Because the LE session necessarily involved light-emitting measurement instruments (Metropsis, retina-WISE, ocular imaging) that were withheld during the DL session to preserve the *<* 10 lux environment, we assessed whether this instrumentation itself measurably perturbed repeated measures of salivary melatonin, subjective sleepiness (KSS), or vigilant attention (PVT), the same outcomes used to establish the LE/DL circadian-phase comparison. Paired Wilcoxon signed-rank tests (with Bonferroni correction for individual time-point comparisons of KSS and PVT, see Code Availability) compared the two sessions across all twelve participants with available data, extending an original seven-participant analysis presented as a poster at SLBTR 2024^16^. Consistent with the original poster findings, melatonin concentrations, subjective sleepiness, and vigilant attention did not differ significantly between LE and DL (p > 0.05 for all comparisons), with only slight, non-significant qualitative trends, shown in Figure 4 (an increase in melatonin and a decrease in vigilant attention, during LE relative to DL).

Verifying the role of a melatonin-specific effect was particularly important because the raw melatonin data were subsequently used for harmonic fitting to extract a circadian period, against which other physiological measures in this dataset were fitted. The light stimuli used to probe retinal function did not significantly perturb the circadian state under the present experimental conditions, supporting the validity of using this FD protocol to characterise non-visual photoreception without confounding downstream effects.

### Completeness verification

Dataset completeness was verified for every participant, session, and modality, by cross-checking the set of files present on the repository against the set expected given the study structure documented in dataset.yaml. The large majority of participant-session combinations are complete. Where files are missing, this often reflects a modality that was simply not scheduled for that session, or isolated device-level failures, such as a detached sensor or a battery or storage limit reached during the 40-hour FD session, rather than the loss of an entire session’s data.

Several modalities (actigraphy, interstitial glucose monitoring, blood pressure and heart rate, skin and core body temperature) involve continuous and long recordings, which are intrinsically challenging and often also require sensors to be switched mid-session. Therefore, each of these has at least one participant-level note describing a data-collection irregularity. Given the modest size of the dataset, these notes are itemised individually rather than aggregated into summary statistics, and are listed in the missing_data.yaml file.

### Known data-quality issues

A number of specific data-quality issues were identified during curation and are documented here and in the accompanying metadata files, so that users can account for them in downstream analyses. Other than this, no reconstruction has been applied to any modality: the released dataset consists exclusively of raw recordings or minimally processed instrument exports, including the documented gaps described in the following list.

- **abpmpro** For participant 209, the PhotoPlethysmoGraphy sensor (PPG) detached during the LE session; the specific set of affected ABPMpro files is documented accordingly.
- **bodycap** During the FD session of participant 210, the third capsule failed and was replaced; therefore, a fourth capsule appears in the 210_bodycap_20240819-21.csv file, which is the one meant to be used in downstream analyses. Moreover, the FD file of participant 211, 211_bodycap_20240718.csv, deviates from the usual BodyCAP raw data scheme: there, the 2nd pill is the one corresponding to the DL session, while the 3rd and the 1st, in this order, measured the FD session.
- **cgm** Two irregularities occurred in continuous glucose monitoring data (participants 202 and 212) and are documented separately in the accompanying cgm_meta_29042025.md metadata file.
- **ibutton** Isolated iButton sensor failures during the FD session led to the loss of specific proximal or distal temperature files for participants 200, 206, and 207. Participants 212 and 213 have a sampling gap of approximately 2 h around the time of the sensor change during the FD session, probably due to accidental detachment of the first sensor from participants’ skin during sleep. Participant 204’s recordings were also affected by some faulty sensors used during the forced desynchrony protocol; in this case, the sensors introduced largely unphysiological random values among the correctly-sampled temperatures. For transparency, the repository includes original raw data; nevertheless, a filtered version of the data and the Python script used to obtain them are available on GitHub^15^ (see Code Availability). All sample timestamps throughout the dataset are recorded in CET/CEST (Central European Time or Central European Summer Time, UTC+01:00/UTC+02:00), with the exception of iButton data, which are sampled in UTC (Coordinated Universal Time). This modality was uploaded to the Edmond repository with its original timestamps. Users are therefore advised to add 1 or 2 h to the timestamps, depending on the time of year, to convert them to CET/CEST before use in analyses, especially if correlating them with other modalities of this dataset.
- **metropsis** For the TCS modality, a subset of files was initially exported under incorrect participant identifiers; these have since been corrected in the released filenames, and the original PDF exports retain a red annotation marking the correction for transparency. For visual stimulation, the dominant eye (determined by the Miles test^17^) was stimulated in all participants to reduce fatigue, while both eyes were tracked throughout; the stimulated eye is treated as primary. One exception occurred for participant 202, whose dominant right eye showed insufficient tracking quality during the first experimental session; consequently, during the forced desynchrony session, the non-dominant left eye was stimulated instead. The identity of each eye is preserved in the corresponding file names.
- **pvt** During the psychomotor vigilance task, in some cases the experimenter had to repeat the measurement when it could not be immediately confirmed that the prior trial had been recorded correctly (e.g. due to a suspected battery issue). As a consequence, duplicate tests were recorded in four cases, producing double the .RTF files: FD session of the pilot and of participants 201 and 202, and LE session of participant 210. As both recordings are valid data and were collected under identical experimental conditions and within a short time window, they constitute two independent, equally valid estimates of the same psychomotor performance. We therefore advise averaging them rather than selecting one file arbitrarily, which would introduce unjustified selection bias, or treating them as independent repeated measures, because both derive from the same participant and session.

## Usage Notes

### Suggested Analyses

Beyond the confirmatory visual-function analyses that motivated data collection, this dataset supports several concrete secondary uses:

- Since core and peripheral temperature (bodycap, ibutton), cardiovascular measures (abpmpro, redcap_logbook), interstitial glucose (cgm), and psychomotor vigilance (pvt) were all recorded during the same 40-hour FD protocol, users can examine within-person coupling across these domains as a function of the imposed sleep-wake schedule and estimated circadian phase, without needing to run a new protocol;
- The pairing of salivary melatonin (saliva) from the DL and FD sessions with the continuous physiological channels makes this dataset suitable for benchmarking or validating circadian-phase estimation algorithms against DLMO-derived phase markers, as already demonstrated for the dlmoR package^12^. The same approach could be extended to phase-estimation from temperature, activity, or cardiovascular signals;
- By using the protocol files as a common timestamp reference across modalities, the dataset offers a realistic testbed for developing or evaluating methods for aligning and resampling heterogeneous physiological time series recorded at markedly different rates and cadences;
- The documented device-level gaps (see Known data-quality issues and Figure 2) provide a real-life setting for developing or benchmarking missing-data handling methods for intensive longitudinal designs, rather than relying on artificially induced missingness.
- Since sleep and wake states were experimentally imposed rather than free-running, the dataset offers the possibility of examining the acute effects of scheduled sleep on any of the recorded modalities. However, users should bear in mind that these states reflect only protocol-derived labels and do not constitute a verified sleep stage.

As an illustrative example of the dataset’s potential, Figure 3 provides the visualisation of a subset of modalities recorded during the FD session for a single participant.

### Use recommendations

To support reuse of the dataset, we summarise here the general and modality-specific caveats that are otherwise documented individually and scattered throughout this article.

First, prior to use, data integrity should be verified against the SHA-256 checksum file included in the repository metadata (SHA256SUMS_v1.0.0.txt), following the instructions provided in the accompanying README.md file.

Timestamps throughout the dataset are expressed in local Europe/Berlin time (CET/CEST, UTC +01:00/+02:00), with the sole exception of the iButton skin-temperature data, which are logged in UTC. To account for the corresponding time offset, these timestamps must be shifted by one or two hours (depending on daylight-saving status) before alignment with any other modality, as was done for Figure 3. This ibutton modality, together with bodycap, additionally presents systematic mid-session sensor handovers, also visible in Figure 3. Rather than treating these data as a single continuous recording, users should explicitly account for the device substitutions by cross-referencing the raw files with the accompanying metadata, since the exact timing of each switch varies by participant.

Another aspect to consider across the entire dataset is that the “Wake” and “Sleep” state labels, wherever applied, reflect the scheduled protocol condition only. No polysomnography was performed, so these labels should not be interpreted as physiologically verified sleep stages.

Furthermore, isolated instrument failures occurred during data collection, and the dataset presents additional modality-specific missingness or irregularities. Minor anomalies were corrected where this could be done without substantially altering the raw state of the data (e.g., mislabeled participant IDs); these corrections are documented in detail in Known data-quality issues. Gaps in the data, by contrast, have not been reconstructed in any form and are provided in their original state; they are documented at the level of individual participant, modality, and session in the missing_data.yaml and other metadata files, as well as in Figure 2. Users should therefore apply a missing-data handling strategy appropriate to their specific research question rather than assume completeness across all participant-modality-session combinations.

To facilitate awareness of these irregularities, and particularly the subtler ones not explicitly visible in Figure 2, the list below summarises all the structural and acquisition-related considerations that are specific to each individual modality.

- **abpmpro** Continuous recordings are segmented according to the imposed sleep/wake schedule using state labels of “Wake”, “Sleep”, or “n/a”; again, these labels reflect the scheduled protocol condition rather than a physiologically verified sleep state. Manual systolic and diastolic blood pressure measurements were taken four times in each experimental block, concurrently with melatonin sampling and prior to each feeding opportunity. Measurements were consistently taken on the right arm, as the left arm was reserved for the ABPMpro cuff, whose measurement was continuous.
- **acttrust** Actigraphy records are organised into three folders corresponding to the three stabilisation phases and named pre-le, pre-fd, and post-fd rather than the usual convention; users should not assume a numeric ordering when parsing folder contents.
- **ambient** One file is provided per room and session combination. File names contain the participant identifier, date, room, and session, allowing files to be matched to the corresponding experimental context directly from the filename.
- **bodycap** Up to four ingestible capsules were administered per participant: three corresponding to the LE, DL, and FD protocol phases, plus an additional capsule administered mid-FD, because a single one could not span the full 40 hours of the protocol. The timing of this second FD capsule varied across participants depending on individual scheduling constraints, and should be confirmed by cross-referencing the raw recordings and metadata, rather than assumed fixed.
- **cgm** This modality was recorded continuously over a 14-day period independent of the experimental session structure. Two independent readers were used, whose cumulative participant data are stored under the group folder. For some participants, data were exported from both readers; users should therefore inspect both exports when working with this modality. Known data irregularities have been identified for participants 202 and 212 and should be inspected before use.
- **eyetracking** Pupil diameter during free viewing was recorded using a Pupil Core eye tracker, and is available for the DC condition only (comprising pilot 113 and participant 202). Recordings were obtained in alternating free-viewing and constrained-viewing blocks. Eye-tracking files can be matched to the corresponding records of Metropsis TCS via the accompanying dc_civibe_TCS-2e_eyetrackin g_blocks.xls file.
- **ibutton** Raw values are logged in UTC+00:00 and must be converted to local CET/CEST before temporal alignment with other modalities. During the FD protocol, two sequential loggers were used per site to accommodate battery and memory limitations; users should account for this handover.
- **metropsis** A subset of participant IDs in the metropsis_tcs submodality were corrected post-hoc following initial data collection. Note that TCS stimulation was delivered monocularly.
- **oct** Left- and right-eye reports are provided as separate files for each session and block. The oct_opticdisc submodality was optional and is therefore available only for a subset of participants.
- **protocol** This file provides a minute-resolution record of the experimental schedule, recapitulating the instructions given to experimenters and participants at each timepoint. Its intended use is limited to cross-referencing timestamps across modalities within a given session, rather than as a primary data source on its own.
- **pvt** In this modality, some duplicate .RTF files exist because experimenters repeated measurements when they suspected recording problems. When device failure in the end did not occur, both records were deemed valid and included in the repository, hence they should be averaged rather than treated as independent observations.
- **redcap** All the variables contained in the questionnaires are defined in the accompanying data dictionaries. Users should note that the “record_id” entry is not standardised to match the participant ID and should only be consulted in case of specific need; otherwise, participant IDs should generally be treated as the informative identifier. Instructions for extracting intraocular pressure, blood pressure, subjective sleepiness and other measurements from redcap/redcap_logbook/group/Civibelogbook_DATA_2024-11-22_1316.csv and redcap/redcap_session_survey/group/Civibesessionsurvey_DATA_2025-08-06_1859.csv are provided in the accompanying data dictionaries: Civibelogbook_DataDictionary_2024-11-22.csv and Civibesessionsurvey_DataDictionary_2025-08-06.csv, respectively.
- **retinawise** Stimulation was delivered monocularly. Data have undergone instrument-internal preprocessing by the acquisition software (see Data processing) but no custom post-processing, and are available for the FD condition only. Users should refer to the associated pupil-response paper^10^ for details regarding the preprocessing steps.
- **saliva** The field “sample_id” encodes the participant, session, and sequential number of each sample. For the FD session of participant 212, the identifier “34” was skipped when assigning sample IDs; the sequence instead runs from 0 to 33 and from 35 to 46, so all 46 physical samples collected for this participant are present.
- **screening** These files include data for participants 203 and 208; both were subsequently excluded from the study and for them no further experimental data are available elsewhere in the dataset.

## Data Availability

The data described in this Data Descriptor^14^ are available at Edmond, the Open Research Data Repository of the Max Planck Society, at DOI: 10.17617/3.JZIJSH, under a Creative Commons Attribution 4.0 (CC BY 4.0) licence. The repository comprises one compressed archive per data modality (listed in the Data Records section above), a METADATA folder with structured .yaml metadata and field-level descriptors, a plain-text file tree, and a SHA-256 checksum manifest.

## Code Availability

No custom code was used to generate the data deposited in the Edmond repository^14^. However, several scripts associated with this Data Descriptor are provided in the following GitHub repository: https://github.com/tscnlab/HeinrichsZanoliThaljiEtAl_bioRxiv_2026.

The repository includes a Python script for filtering non-physiological temperature measurements recorded by a faulty iButton sensor (see Known data-quality issues), together with the corresponding output files, named .FILTERED.xlsx. In addition, a second Python script is provided to reproduce Figure 3 of this article, along with three REDCap-exported datasets (blood pressure, intraocular pressure, subjective sleepiness) and the participant-209-specific glucose dataset exported from the group-level file; these datasets are ready for use and may otherwise be difficult to identify within the complete dataset. The figure was subsequently adjusted manually, introducing a break in the y-axis of the core body temperature subplot. Likewise, Figure 1 and Figure 2 were produced manually and therefore no code is available to reproduce them. Finally, the GitHub repository contains the complete raw data sources needed to reproduce the figures and analyses, the R scripts, and the results of the paired Wilcoxon tests used to verify the absence of circadian effects associated with the use of light-emitting devices.

## Author Contributions

H.S.H.: Conceptualization, Methodology, Investigation, Data curation, Project administration, Writing – original draft, Writing – review & editing. C.Z.: Data curation, Visualization, Writing – original draft, Writing – review & editing. S.M.T.: Investigation, Data curation, Writing – review & editing. L.G.: Investigation, Writing – review & editing. E.K.: Investigation, Writing – review & editing. M.S.: Conceptualization, Methodology, Resources, Supervision, Project administration, Funding acquisition, Writing – review & editing.

## Acknowledgements

We thank the fine mechanical workshop of the Max Planck Institute for Biological Cybernetics for constructing the sleep laboratory used in this study, Jannik Romanowski and Johannes Boldt for customising the experimental setup, Martial Geiser for designing the optical components and supporting the device engineering, Joachim Werner for providing emergency IT support and rapid troubleshooting at short notice throughout the data collection, Katharina Stein and Constanza Bravo for helping to pilot the protocol and for informing the refinement of the experimental procedure, and all participants for their sustained commitment across the weeks-long protocol.

